# Zirconium(IV)-IMAC for phosphopeptide enrichment in phosphoproteomics

**DOI:** 10.1101/2020.04.13.038810

**Authors:** Ignacio Arribas Diez, Ireshyn Govender, Previn Naicker, Stoyan Stoychev, Justin Jordaan, Ole N. Jensen

## Abstract

Phosphopeptide enrichment is an essential step in large-scale, quantitative phosphoproteomics studies by mass spectrometry. Several phosphopeptide affinity enrichment techniques exist, such as Immobilized Metal ion Affinity Chromatography (IMAC) and Metal Oxide Affinity Chromatography (MOAC). We compared Zirconium (IV) IMAC (Zr-IMAC) magnetic microparticles to more commonly used Titanium (IV) IMAC (Ti-IMAC) and TiO_2_ magnetic microparticles for phosphopeptide enrichment from simple and complex protein samples prior phosphopeptide sequencing and characterization by mass spectrometry (LC-MS/MS). We optimized sample-loading conditions to increase phosphopeptide recovery for Zr-IMAC, Ti-IMAC and TiO_2_ based workflows. The performance of Zr-IMAC was enhanced by 19-22% to recover up to 5173 phosphopeptides from 200 µg of protein extract from HepG2/C3A cells, making Zr-IMAC the preferred method for phosphopeptide enrichment in this study. Ti-IMAC and TiO_2_ performance were also optimized to improve phosphopeptide numbers by 28% and 35%, respectively. Furthermore, Zr-IMAC based phosphoproteomics in the magnetic microsphere format identified 23% more phosphopeptides than HPLC-based Fe(III)-IMAC for same sample amount (200 µg), thereby adding 37% more uniquely identified phosphopeptides. We conclude that Zr-IMAC improves phosphoproteome coverage and recommend that this affinity enrichment method should be more widely used in biological and biomedical studies of cell signalling and in the search for disease-biomarkers.

## INTRODUCTION

Reversible protein phosphorylation regulates multiple processes in the cell, such as cell differentiation, metabolism, apoptosis and growth[1–3]. The reversible protein phosphorylation process is controlled by protein kinases that add the phosphate group to e.g. serine, threonine or tyrosine amino acid residues, and protein phosphatases that remove phosphate groups from amino acid side chains[3]. Alterations or dysregulation of these protein phosphorylation mechanisms can cause severe diseases such as cancer [4–6], diabetes[7, 8] and neuronal disorders[9, 10]. Therefore, therapeutics that target phosphoproteins and protein kinases have become important tools for battling these illnesses[11, 12].

Mass spectrometry (MS) is a key technology for defining the proteome, identifying new biomarkers and understanding cellular and molecular regulatory mechanisms. MS is the preferred bioanalytical method for in-depth analysis of post-translational modifications of proteins [13], and it is indispensable for profiling and quantifying the dynamic phosphoproteome [14–16]. MS based phosphoproteomics strategies rely on efficient enrichment of phosphopeptides (p-pep) since transient protein phosphorylation events frequently occur at low stoichiometry [14, 17]. Commonly used p-pep affinity enrichment techniques include anti-pTyr/pSer/pThr antibodies [18–20], pTyr-superbinders [21], immobilized metal ion affinity chromatography (IMAC) and metal oxide affinity chromatography (MOAC). IMAC uses metal cations, and MOAC metal oxides, that interact with the oxygen atoms of phosphate groups in peptides [17, 22, 23]. More recently, molecularly imprinted polymers (MIPs) have emerged as potential tools for selective phosphopeptide enrichment [24].

IMAC often employs Fe^3+^ as the phosphate-chelating ion species [25], but Ga^3+^, Ti^4+^ and Zr^4+^ are viable alternatives [26–29]. Particularly, Ti^4+^ has gained increased utility in recent years, i.e. Ti-IMAC [30–32]. TiO_2_ is the preferred reagent for MOAC based methods [33, 34], but other metal oxides have been tested, including Al_2_O_3_, Fe_2_O_3_ and ZrO_2_ [35–37]. Previous studies compared the efficiency of Fe-IMAC, Ti-IMAC, TiO_2_ and ZrO_2_ for phosphopeptide enrichment [38–41], demonstrating complementarity between these approaches and that their combination may increase the coverage of the phosphoproteome.

Zirconium was first employed for phosphoproteomics in 2006 as ZrO_2_ [37] and soon after as Zr^4+^ phosphonate for enrichment and MALDI MS analysis of phosphopeptides [42]. A year later, Feng et al. published the first off-line Zr-IMAC method for p-pep LC-MS/MS analysis [26]. Since then, Zirconium has been utilised in different formats, such as magnetic core silica nanoparticles [43], monolith columns[44], zirconium arsenate-based materials [35, 45] and aerogels [46]. Studies of Zr based techniques focused on ZrO_2_ and their performance were usually compared to TiO_2_ [37, 47–49]. Furthermore, Zr-IMAC materials were compared to ZrO_2_ or Fe-IMAC [26, 43], showing higher selectivity than Fe-IMAC and improved stability against salt-containing solvents in the context of strong cation exchange (SCX) chromatography separation of peptides. Despite these promising reports, Zr-based affinity enrichment techniques have been overshadowed by the widely used TiO_2_ and Ti-IMAC approaches in phosphoproteomics [31, 50].

Zirconium is a transition metal from group 4 in the periodic table, i.e. the same group as Titanium. Thus, Zr has very similar properties as Ti, but with distinct differences. Both Zr and Ti can oxidize to state four (IV), producing salts of ZrO_2_ and TiO_2_ that strongly interact with oxygen anions. Zr is heavier than Ti (91.22 Da vs. 47.86 Da), making it a stronger Lewis acid, i.e. electron-pair acceptor, with a higher coordination number in the crystalline form (7 for ZrO2 and 6 for TiO_2_) [37, 51]. Since they are high-valence metal cations, both Zr and Ti have a unique coordination specificity with phosphates that allow binding of more than one phosphate group [52–54], making them both excellent candidates for phosphopeptide enrichment.

To our knowledge, a detailed comprehensive assessment of Zr-IMAC for phosphopeptide enrichment using complex biological samples is lacking. It remains to be established whether Zr-IMAC is a robust and efficient method for mass spectrometry based phosphoproteomics studies.

We hypothesized that Zr (IV) can perform similarly to Ti (IV) for phosphopeptide enrichment using polymer-based IMAC magnetic microparticles [31]. We also hypothesized that optimization of the protocols and reagents may improve performance for each of these methods, albeit not with the same specific conditions.

We therefore sought to improve phosphopeptide enrichment efficiency by adding hydroxy acids to the sample loading solvent, as hydroxy acids are known to improve the specificity and selectivity of phosphopeptide enrichment using IMAC and MOAC [48, 55]. We evaluated the effects of glycolic (GA), lactic (LA) and tartaric (TA) acids on p-pep enrichment using Zr-IMAC, Ti-IMAC and TiO_2_ magnetic microparticles. Our optimized Zr-IMAC protocol selectively and efficiently captures p-pep from microgram amounts of protein starting material obtained from biological samples.

## EXPERIMENTAL PROCEDURES

### Experimental design and statistical rationale

A total of 36 digest (20 µg each) of BSA + Casein were used for the optimization of binding solvent composition. Six separate enrichments were performed in technical duplicates with a different binding solvent (Std, S1, S2, S3, S4, S5, see Table 1) with three types of magnetic microparticles (Zr-IMAC, Ti-IMAC and TiO_2_). The standard binding solvent (Std) served as a control. Zr-IMAC, Ti-IMAC and TiO_2_ magnetic microparticles were evaluated in n = 3 technical replicates for phosphopeptide enrichment in two experimental binding solvent conditions: standard (Std) as the control and optimized (S1/ TiO_2_, S2/Zr-IMAC, S3/Ti-IMAC). In total, 36 samples of 200 µg of HepG2/C3A human hepatocyte cells tryptic digest were used. For the comparison of optimized microparticles with Fe-IMAC HPLC, n = 4 technical replicates of 200 µg of HepG2/C3A human hepatocyte cells tryptic digest were enriched for n = 5 experimental conditions (total of 20 samples). Statistical tests used are described within each section.

**Table 1.** List of solvents used during phosphopeptide enrichment with magnetic microparticles.

| Solvent name | Enrichment step | Binding solvent composition |
| --- | --- | --- |
| Std | Binding/Washing | 80% MeCN, 5% TFA, 1M GA |
| S1 | Binding/Washing | 50% MeCN, 0.1% Acetic, 0.1M GA |
| S2 | Binding/Washing | 80% MeCN, 5% TFA, 0.1m GA |
| S3 | Binding/Washing | 80% MeCN, 5% TFA, 0.1M LA |
| S4 | Binding/Washing | 80% MeCN, 5% TFA, 0.1M TA |
| S5 | Binding/Washing | 50% MeCN, 0.1% Acetic, 0.1M TA |
| Wash 1 | Washing | 80% MeCN, 1% TFA |
| Wash 2 | Washing | 10% MeCN, 0.2% TFA |
| Elution | Elution | 1.25M Ammonium hydroxide |

### Materials and Chemicals

All chemicals were obtained from Sigma Aldrich (Merck KGaA) unless otherwise stated. Ti(IV)-IMAC, Zr(IV)-IMAC, and TiO_2_ ferromagnetic microparticles for phosphopeptide enrichment were from ReSyn Biosciences, RSA (Cat. no. MR-TIM002, MR-ZRM002 and MR-TID002). Kingfisher deep-well 96 plates were purchased from ThermoFisher Scientific. Sep-Pak tC18 3 cc Vacuum Cartridges were purchased from Waters Corp (Cat. no. WAT054925).

### Cell culture and lysis

HepG2/C3A human hepatocyte cells (ATCC CRL-10741) were kindly provided by associate professor Adelina Rogowska-Wrzesinska and Helle Frandsen (University of Southern Denmark, Odense, DK). Cells were in standard culture conditions (87.5% D-MEM (containing 1 g glucose/L) (Gibco), 1% Non-Essential Amino Acids (Gibco), 10% FCS (Foetal calf serum), 0.5% Penicillin/Streptomycin (Gibco), 1%GlutaMAX (Gibco), 37°C, 5% CO2 95% air) and, when needed, they were trypsinized for 3 min with 0.05% Trypsin/EDTA (Gibco Cat. no. 15400-054), diluted 1:4 and sown out into falcon tubes or microtitre plates [56]. Cells were collected 5 days after trypsinization at 85% confluence at subculture 10 and washed 5 times with warm (37°C) Hanks’ Balanced Salt Solution (HBSS without Ca++ and Mg++, Gibco Cat. no. 14175-053). After removal of all the remaining HBSS, the samples were snap frozen in liquid nitrogen and stored at −80°C until further processing.

Frozen cell pellets were thawed on ice and suspended with gentle pipetting in 5 mL of cold lysis buffer (1% sodium deoxycholate (SDC), 100mM ammonium bicarbonate (ABC), 10mM Tris(2-carboxyethyl)phosphine hydrochloride (TCEP), 40mM 2-Chloroacetamide (CAA), pH 8.5, one tablet per 20mL cOmplete^TM^ Protease Inhibitor Cocktail (Roche), 1 tablet per 10mL PhosSTOP^TM^ (Roche)), then ruptured with a probe sonicator model CL-18 (Qsonica) in an ice-bath for 5 cycles of 20 sec 50% power and 20 sec break. The liquid was then heated at 80ºC for 10 min. Proteins were precipitated with acetone overnight and washed twice with ice cold 80% acetone. The pellet was re-suspended in 5mL of 1% SDC + 50mM ABC pH 8.5 and protein concentration was measured with Pierce BCA protein assay kit (Thermo Scientific). Samples were then aliquoted into 1.5mL LoBind eppendorf tubes (Sorenson BioScience) and stored at −80ºC for future use.

### Tryptic digestion of (phospho)protein samples

For the phosphopetide binding/elution solvent optimization experiments, a 5 mg mixture of α-, β- and κ-Casein (Cat. no. C7078) and 5 mg of Bovine Serum Albumin (BSA, Hyclone, Cat. no. SHB0574) were dissolved in 150 µl and 200µl, respectively, resuspension buffer (5.5M Guanidinium-HCl in 50mM Ammonium Bicarbonate (ABC), 50mM DTT) and heated at 35°C for 10 min. Ten microliters of BSA solution were mixed with 30uL of Casein solution to a final concentration of 0.45mg/mL and incubated at 60°C for 1h for reduction, followed by *S*-alkylation with 10uL of 0.5M IAA in 50mM ABC and incubated at room temperature in the dark for 30min. The protein mixture was diluted to 25mg/ml with 50mM ABC and 500ug of protein were digested with 2% w/w Trypsin (Promega, Cat. no. V5111) in 1mL of 50mM ABC overnight at 37°C. Digestion was stopped with 10% formic acid (FA) to a final concentration of 0.5% FA and 45uL aliquots were stored at −20°C.

Cell lysate protein extracts were digested with in-house methylated [57] porcine Trypsin (Cat. no. T0303) at 2% w/w and incubated overnight at 37°C. Tryptic digestion was terminated by adding trifluoroacetic acid (TFA) to 1% concentration. Digested samples were centrifuged for 5 min at 14,000 rpm to precipitate the SDC and the supernatant liquid was transferred to a new tube. Peptide samples were then de-salted using Sep-Pak tC18 3 cc Vac Cartridges (Waters Corp, Cat. no. WAT054925). Cartridges were first conditioned with 2mL of 95% acetonitrile (MeCN) / 0.1% TFA and equilibrated with 2x 3mL 5% MeCN / 0.1% TFA. Peptide samples were then loaded and the cartridge was rinsed using 2x 3mL of 5% MeCN/0.1% TFA. Elution was achieved by adding 3x 1mL 50% MeCN/0.1% TFA. Eluted peptide samples were collected in 5mL tubes. Peptide samples were dried using a vacuum centrifuge concentrator and stored at −80°C.

### Phosphopeptide enrichment using polymer based magnetic microparticles

Optimization of sample loading conditions: TiO_2_, Ti-IMAC and Zr-IMAC magnetic microparticles (ReSyn Biosciences) were tested in duplicate with the standard loading and binding solvent recommended by the manufacturers (80% MeCN, 5% TFA, 1M glycolic acid (GA)) and five alternative solutions: S1 (50% MeCN, 0.1% acetic acid (AcA), 0.1M GA), S2 (80% MeCN, 5% TFA, 0.1M GA), S3 (80% MeCN, 5% TFA, 0.1M lactic acid (LA)), S4 (80% MeCN, 5% TFA, 0.1M tartaric acid (TA)) and S5 (50% MeCN, 0.1% AcA, 0.1M TA).

Dried tryptic peptides were re-dissolved in 0.1% TFA to a concentration of ~4 µg/µL and diluted to a final volume of 200uL with the corresponding binding solvent (S1-S5) for enrichment. Experiments performed at Council for Scientific and Industrial Research (CSIR, Pretoria, Republic of South Africa) employed a KingFisher Duo workstation (Thermo Scientific) for automated magnetic microsphere based phosphopeptide enrichment. The protocol was adapted for manual magnetic microsphere-based p-pep enrichment at the University of Southern Denmark (Odense, Denmark). The enrichment protocol was performed as previously described [31], maintaining a 1:2 peptide-to-beads weight ratio. In brief, the magnetic particles were first equilibrated with loading/binding solvent for 5 min, followed by 20 min incubation with the peptide mixture (20 µg for Casein/BSA tryptic digests, 200 µg for liver cell protein extracts). Microspheres were then washed for 2 min each, first with solvent (S1-S5, 500 µL), then wash solvent 1 (80% MeCN + 1% TFA, 500 µL) and lastly by wash solvent 2 (10% MeCN + 0.2% TFA, 500 µL). Phosphopeptides were eluted from the microparticles by 10 min incubation in 1.25M of ammonium hydroxide solution (200 µL). Total program time was 45 min and all incubation steps were done with continuous mixing to keep a homogenous bead suspension. Collected peptides were lyophilized and stored at −80ºC prior analysis.

For Zr-IMAC reproducibility experiments, Zr-IMAC and Ti-IMAC magnetic microbeads were tested in five replicates with the standard binding solvent and the optimized solvent S2. Enrichment protocol [31] was manually performed on a 96×2 mL plate. A schematic visualization of the workflow can be seen in Figure 1B, right branch.

**Figure 1.**
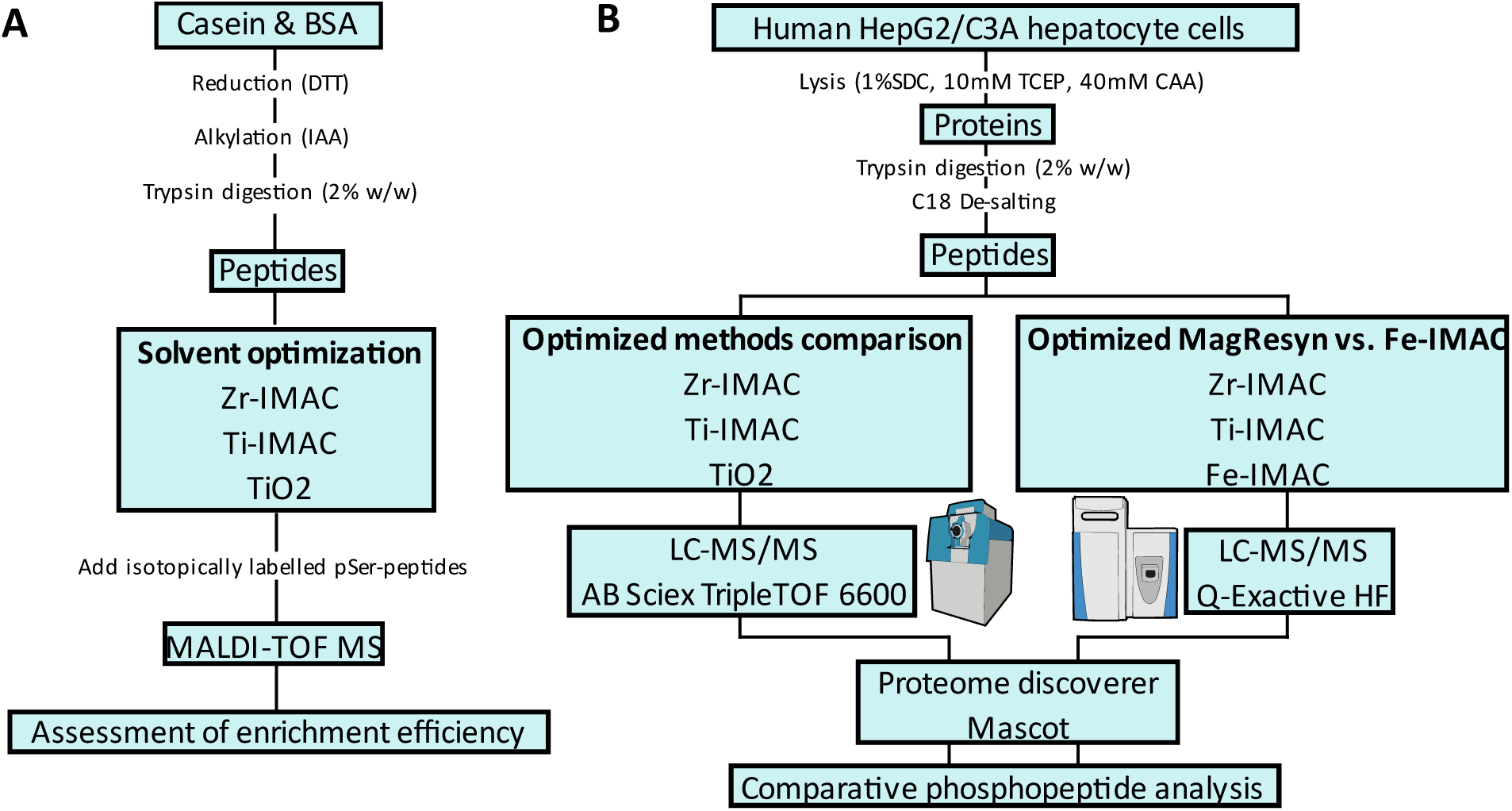
Schematic description of the workflows. (A) Selection of enrichment solvent conditions for each bead chemistry using simple peptide digests and spiking them with three isotopically labelled peptides for MS1 relative quantification of phosphopeptides. (B) Test of optimal conditions against complex peptide mixtures. First, the respective optimal solvents for Zr-IMAC, Ti-IMAC and TiO_2_ were evaluated (left branch). The best performing solvent and microparticles were afterwards compared with Fe-IMAC HPLC enrichment (right branch).

### Phosphopeptide enrichment with Fe-IMAC HPLC

LC based Fe(III)-IMAC [38] was implemented using a ProPac^TM^ Fe-NTA column (2×50 mm, Thermo Scientific) connected to a UPLC Ultimate 3000 system (Thermo Scientific) via an 800 µL injection loop. The IMAC column was charged with iron (III) according to the manufacturer’s instructions. Prior sample loading the column was equilibrated with 50% solvent C (100% MeCN) + 50% solvent A (99.9% H2O + 0.1% TFA). Approximately 200 µg of tryptic protein digest was dissolved in 0.1% TFA to 4ug/µL concentration and diluted to 710uL 50% MeCN + 0.1% TFA. Injection volume was 700 µL. After sample loading, the column was washed for 5 min with 50% solvent C + 50% solvent A followed by 50% solvent C + 50% solvent B (20mM NH4OH) for 2 min. Phosphopeptides were collected with 100% solvent B for 2 min and the column was re-equilibrated with 50% solvent C + 50% solvent A for 7 min. Peptides were dried using a vacuum centrifugal concentrator and stored at −80°C prior LC-MS/MS analysis.

### MALDI-TOF MS analysis of peptides

Phosphopeptide samples obtained by enrichment from Casein/BSA tryptic digests were re-suspended in 25uL of 0.1% TFA. Two microliters of the peptide solution were mixed with 2uL of a 1:3:3 mixture of isotopically labelled p-pep (VPQ[L_C13N15]EI[V_C13N15]PN[pSer]AEER (1662 Da), YKVPQLE[I_C13N15]VPN[pSer]AEER (1957 Da) and FQ[pSer]EEQQQTEDE[L_C13N15]QDK (2067 Da) (ThermoFisher Scientific, Supplementary Figure 1)) and 2uL of matrix solution (30 mg/mL DHB in 50% MeCN + 1% H3PO4), plated 1.8uL on a 384 ground-steel MALDI plate and once dried, re-crystallized with 0-5uL acetone.

Peptides were analysed on a Bruker Autoflex MALDI-TOF MS instrument using positive ion reflector mode and a m/z range of 1000-3500. Pulsed ion extraction was 40ms and a UV laser power of 30-45% were applied. Spectra corresponding to three hundred laser pulses were accumulated per acquisition. The signal for each p-pep was normalised against the highest responding heavy internal standard (YKVPQLE[I_C13N15]VPN[pSer]AEER, m/z 1958) to quantify enrichment efficiency for each solvent composition. Experiments were analysed in duplicate.

### Phosphopeptide analysis by LC-MS/MS using Sciex TripleTOF 6600 system

Dried phosphopeptides were dissolved in 10uL of 2% MeCN + 0.1% FA, vortexed for 1 min and centrifuged at 14,000 rpm for 10 min to remove any residual particles. Peptides were analysed using a Dionex Ultimate 3000 RSLC system (Thermo Scientific) coupled online to a TripleTOF® 6600 hybrid tandem mass spectrometer (Sciex). Injected peptide samples were on-line de-salted/concentrated using an Acclaim PepMap C18 trap column (75 μm × 2 cm, Thermo Scientific) for 2 min at 5uL/min using 2% MeCN + 0.2% FA. Trapped peptides were gradient eluted and separated on an Acclaim^TM^ PepMap^TM^ C18 RSLC column (75 μm × 15 cm, 2 µm particle size, Thermo Scientific) at a flow-rate of 500nL/min with a linear gradient of 3%-40% solvent B (80% MeCN + 0.1% FA) against solvent A (100% H2O + 0.1% FA). Eluted peptides were electrosprayed using a NanoSpray® III Ion Source (Sciex) with the following source settings: CUR - 10, GS1 - 24, GS2 - 0, HT - 80° C and ISVF - 2800.

LC-MS/MS was performed in the data-dependent acquisition (DDA) mode. Mass spectra were acquired in the range *m/z* 400-1500 (2+ to 5+ charge states) using an accumulation time of 250 ms followed by 80 MS/MS events. MS/MS spectra were acquired for the range m/z 100-1800 at 25 ms accumulation time per spectrum. Minimum threshold for triggering MS/MS was set to 250 cps. Precursor ions were fragmented in the q2 collision cell using nitrogen as the collision gas. Collision energies were automatically defined as function of m/z and charge state. Selected ions for MS/MS were dynamically excluded for 15 sec.

### Phosphopeptide analysis by LC-MS/MS using Q-Exactive HF system

Dried phosphopeptides were handled as previously mentioned and analysed on a Dionex Ultimate 3000 RSLCnano system coupled online to a Q-Exactive® HF hybrid quadrupole-orbitrap tandem mass spectrometer (Thermo Scientific). Injected peptides were on-line de-salted using a C18 PepMap^TM^ 100 trap column (300um i.d. x 5mm, 5um, 100Å, Thermo Scientific) for 2 min using 2% MeCN + 0,1% FA. Trapped peptides were gradient eluted and separated on a 75um i.d. x 30cm column packed inhouse with ReproSil-Pur 120 C18 1.9um particles (Dr. Maisch) at a flow-rate of 300nL/min with a linear gradient of 5%-35% solvent B (95% MeCN + 0.1% FA) against solvent A (100% H20 + 0.1% FA).

Precursor ion (MS) spectra were acquired in the range m/z 300 - 2000 (2+ to 5+ ion charge state) at a mass resolution setting of 60,000 at m/z 400 in profile mode, with an AGC target of 1e5 ions. The 20 most intense precursor ions were fragmented with higher-energy collisional dissociation (HCD) and dynamically excluded for 15 sec. Fragments were detected in the Orbitrap at a mass resolution setting of 30,000 at m/z 400 in centroid mode.

### LC-MS/MS Data analysis

Prior data analysis, all wiff data files from Sciex TripleTOF® 6600 were converted to Mascot Generic File (mgf) with ProteoWizard MSConvert (version 3.0.19137) with peak picking filtering for the 150 most intense peaks. Mgf files and raw files from the Q-Exactive® HF were processed with Proteome Discoverer 2.3 (Thermo Scientific) and searched against SwissProt human database (February 2019 release) using Mascot 2.6 (Matrix Science) with 50 ppm precursor mass tolerance for mgf files and 10 ppm for raw files. Fragment mass tolerance was set to 0.2 Da for mgf files and 0.05 Da for raw files with a limit of 2 missed tryptic cleavages. *S*-carbamidomethylation of cysteine was set as a fixed modification while *O*-phosphorylation of serine, threonine, and tyrosine, and oxidation of methionine were set as variable modifications. Peptides were filtered using a Mascot ion score threshold of ≥ 18 and a FDR of ≤ 0.01 through Percolator [58]. Phosphorylation localization probabilities were calculated with ptmRS [59] and only probabilities ≥ 75% were accepted. Feature analysis of phosphopeptides was done using a custom made R script to calculate peptide hydrophobicity by Kyte & Doolittle hydropathy indexes [60] and the pI values were obtained from the online protein isoelectric point calculator [61]. An Excel sheet was used to count the number of basic and acidic residues, as well as the total number of amino acids per sequence. Protein sequence visualization and phosphorylation site illustration was done on the General Protein/Mass Analysis for Windows software tool (GPMAW v. 11.0)[62].

The mass spectrometry proteomics data described in this manuscript have been deposited to the ProteomeXchange Consortium (http://www.proteomexchange.org) via the PRIDE partner repository [63] with the dataset identifier PXD018273.

## RESULTS

### Optimization of solvent composition for phosphopeptide enrichment by IMAC and MOAC

Three magnetic bead chemistries (two IMAC, one MOAC) were tested with recommended solvents (Std) and five modified binding solvents (S1-S5) to identify the optimal bead-solvent combination (Table 1). Tryptic digests of Casein/BSA were used for these initial phosphopeptide enrichment experiments (Figure 1A). Zr-IMAC, Ti-IMAC and TiO_2_ microbeads were incubated in their corresponding binding solvents with the protein digests, followed by washing of the beads and elution of the p-pep with 1.25 M ammonium hydroxide solution.

To quantify phosphopeptide recovery, the eluted fractions were mixed with three isotopically labelled phosphopeptides (Supplementary Figure 1) prior MALDI analysis, and the light phosphopeptide intensities were normalized to the most intense signal of the heavy labelled peptides (YKVPQLE[I_C13N15]VPN[pSer]AEER, m/z 1958). Figure 2 shows the relative intensities of identified phosphorylated sequences. In all cases, the addition of tartaric acid to the sample loading solvent (solvents S4 and S5, Figure 2) resulted in a significant decrease in phosphopeptide signal intensity in MALDI MS. Reducing the glycolic acid concentration from 1 M to 0.1M (S2) enhanced the phosphopeptide signal intensity without major effects on specificity (Std vs S2, Figure 2). Also, 0.1M lactic acid (S3) improved phosphopeptide signal intensities without major effects on specificity (Std vs S3, Figure 2). For Ti-IMAC and Zr-IMAC, the very acidic solvents S2 and S3 (5% trifluoroacetic acid) performed better than solvent S1 (0.1% Acetic acid) as seen by the higher intensity values obtained from most phosphopeptides (Figure 2). However, TiO_2_ showed augmented signal intensity when using solvent S1 at milder pH conditions (S1 vs. S2 and S3, Figure 2).

**Figure 2.**
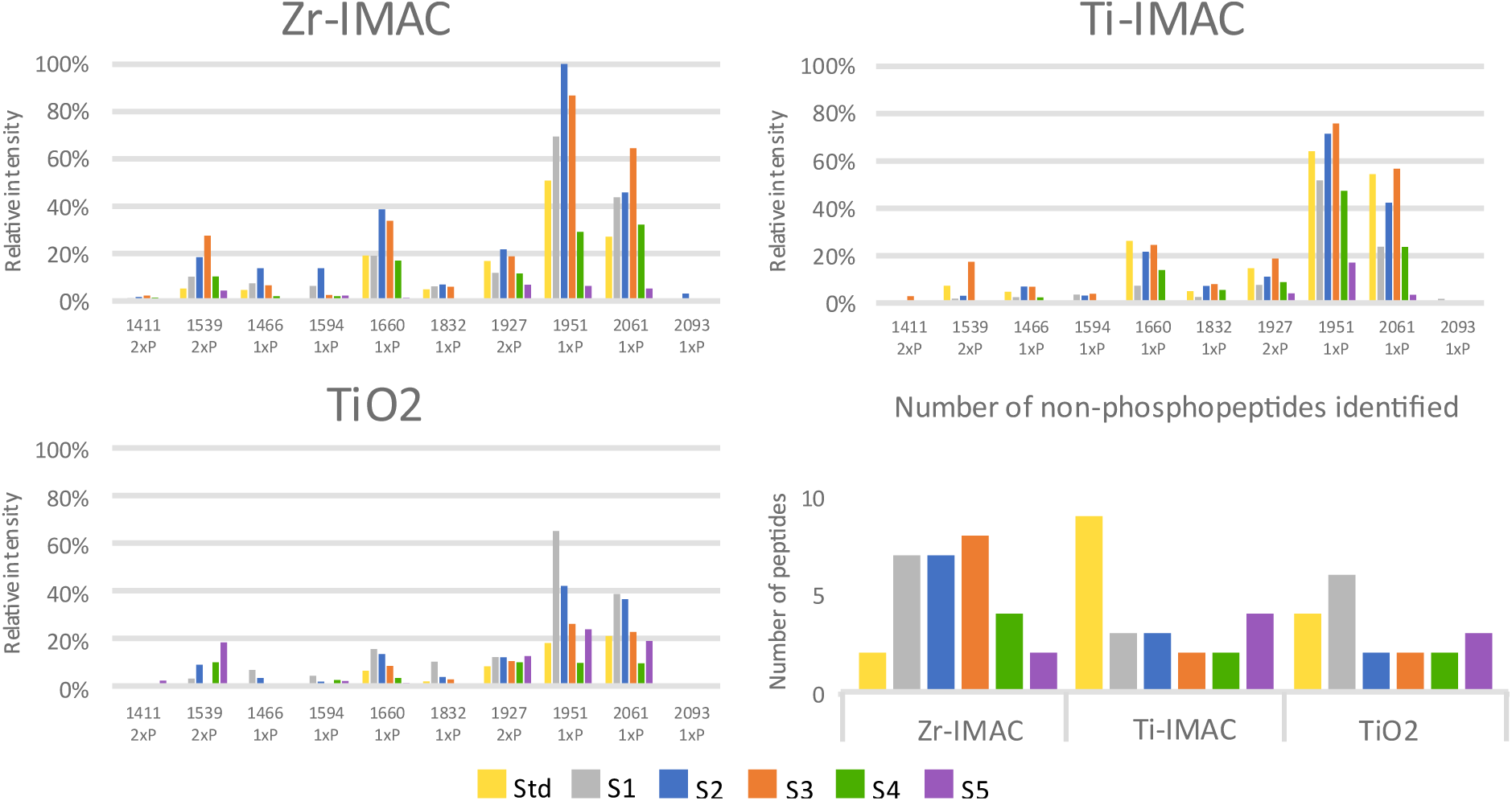
Phosphopeptide enrichment efficiency of each condition based on the relative intensity of the unlabelled phosphopeptides vs. the most intense stable isotope labelled phosphopeptide (m/z 1958). Bottom right, number of non-phosphorylated peptides identified for each condition (Refer to table 1 to see solvent compositions).

Relative phosphopeptide signal abundances were higher with S2/Zr-IMAC than with Std solvent (Figure 2), doubling the intensities of p-pep m/z 1660 and m/z 1951 and tripling the signal intensity at m/z 1539 (Figure 2, and data not shown). S3/Ti-IMAC and S1/ TiO_2_ showed similar results when compared to the Std solvent: S1/ TiO_2_ improved by 2x, 3x and 5x the ion signals of p-pep at m/z 1660, 1951, 1832, respectively, while S3/Ti-IMAC doubled that of m/z 1539.

Based on these results, solvents S3, S2 and S1 were selected as optimal loading solvents for Ti-IMAC, Zr-IMAC and TiO_2_, respectively, and further tested by using complex peptide mixtures derived from cell extracts.

### Enrichment efficiency assessment using complex peptide mixtures

Zr-IMAC, Ti-IMAC and TiO_2_ functionalized magnetic microparticles were tested for phosphopeptide enrichment using the standard and optimized solvent conditions. A complex sample consisting of a tryptic peptide mixture derived from human HepG2/C3A cells was used for automated phosphopeptide enrichment [31] on a KingFisher Duo workstation (Thermo Scientific). The enriched phosphopeptide eluates were analysed by LC-MS/MS on a Sciex TripleTOF 6600 and the raw files were processed for peptide identification using Proteome Discoverer and Mascot as the search engine (Figure 1B, left branch).

Changing the standard binding solvent to an optimized binding solvent (S1-S3) increased the number of identified phosphopeptides for all bead types (Figure 3A). Zr-IMAC retrieved more p-pep than Ti-IMAC and TiO_2_ in both standard (3900) and optimized (4624) conditions, respectively (Figure 3A). Moreover, S2/Zr-IMAC captured the highest number of p-pep (4624) followed by S3/Ti-IMAC (4042). S1/ TiO_2_ recovered a higher number of p-pep (2744) as compared to using the Std solvent (2039), albeit lower numbers than obtained by either Zr-IMAC or Ti-IMAC. Based on enumeration of retrieved and identified phosphopeptides, Zr-IMAC and Ti-IMAC outperformed TiO_2_ under standard conditions and optimized conditions, identifying almost twice the number of p-pep under Std solvent conditions (Figure 3A).

**Figure 3.**
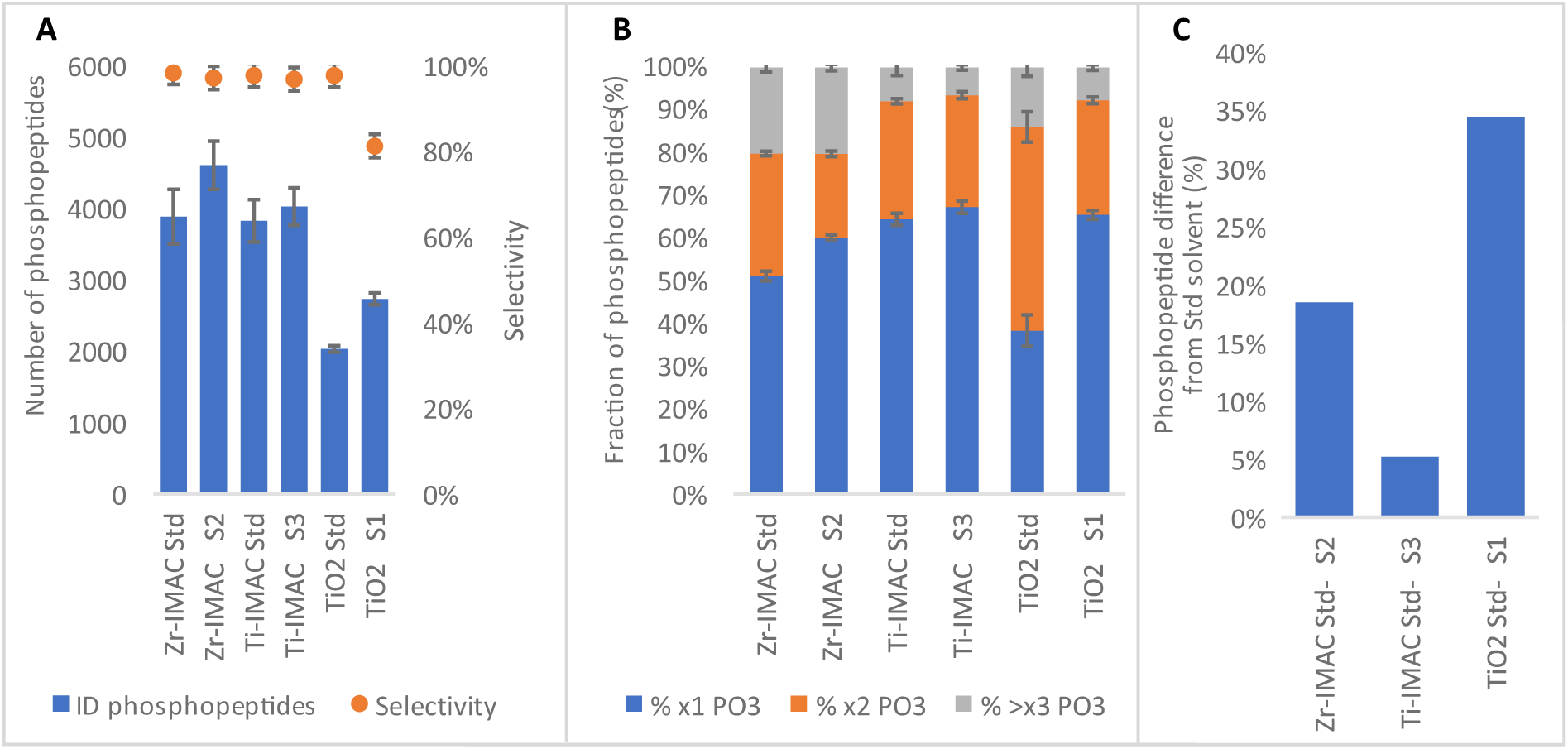
Phosphopeptide (p-pep) profiling of MagReSyn® microparticles by LC-MS/MS on AB Sciex TripleTOF 6600. (A) Number of p-pep identified for each condition (blue bars) and the selectivity of each enrichment (orange dots). (B) Mono-(blue), di-(orange) and multi-phosphorylated (grey) peptide distribution in each condition. (C) Difference in percentage of identified p-pep between the optimized solvent (S1, S2, S3) and the standard (Std).

Next, we investigated the selectivity of each enrichment method for phosphopeptide enrichment (Figure 3A). Zr-IMAC and Ti-IMAC showed over 97% selectivity for p-pep as determined by the ratio: (number of identified p-pep)

/ (number of identified peptides + p-pep), while TiO_2_ selectivity was reduced to 82% when using S1 binding solvent (0.1% Acetic acid) as compared to the Std control (98%). This was to be expected, as the acidic environment in S1 was less harsh than Std binding solvent, allowing un-specific binding of carboxylate groups to the metal oxide of TiO_2_.

Notably, the optimized binding solvents altered the relative distribution of singly, doubly and multiply phosphorylated peptides for all three affinity enrichment chemistries, increasing the number of mono-phosphorylated peptides relative to di- and multi-phosphorylated peptides (Figure 3B). This effect was most pronounced for TiO_2_, and smaller for Zr-IMAC and Ti-IMAC.

The overall improvement of phosphopeptide recovery by tailored and optimized binding solvent is shown in Figure 3C. The TiO_2_ method gained the most from the optimization, binding 35% more p-pep when using S1 as compared to Std conditions, followed by S2/Zr-IMAC (19% improvement) and S3/Ti-IMAC (5% improvement).

Considering the protein level, optimized S2/Zr-IMAC protocol identified 1733 phosphoproteins, thereby exceeding all other methods and conditions that ranged from 930 to 1545 phosphoproteins (Supplementary Figure 2). Ti-IMAC identified a similar number of phosphorylated proteins (1499 and 1478, respectively) and sharing 85% of the total proteins with Zr-IMAC. TiO_2_ identified the lowest number of phosphoproteins but gained the most proteins (34%) from a change of solvent condition from Standard to S1. Ti-IMAC improved by 5% more proteins by using solvent S3 rather than Std. Using the PANTHER GO-slim tool for genome classification [64] we analysed the identified proteins for the optimized methods on their cellular component to observe their proteome coverage (Supplementary Figure 3). All three materials presented the same distribution of protein classes: around 80% representing cell (GO:0005623) and organelle (GO:0043226), followed by ~15% of protein-containing complexes (GO:0032991) and ~3% membrane proteins.

In total, 5173 phosphorylated peptides were identified by the combination of S2/Zr-IMAC, S3/Ti-IMAC and S1/ TiO_2_ (Figure 4A). Almost half of these phosphopeptides (44%) were recovered by all three methods (2278, Figure 4A). Some phosphopeptides were exclusively recovered by one method (S2/Zr-IMAC: 829/16%; S3/Ti-IMAC: 270/5.2%; S1/ TiO_2_: 85/1.6%; Figure 4A), which demonstrated that Zr-IMAC contributes a significant number of additional phoshopeptides as compared to other methods. Furthermore, S2/Zr-IMAC recovered 89.3% of the total number of phosphopeptides (Figure 4A) identified by all methods, followed by S3/Ti-IMAC (79.7%) and TiO_2_ (53%).

**Figure 4.**
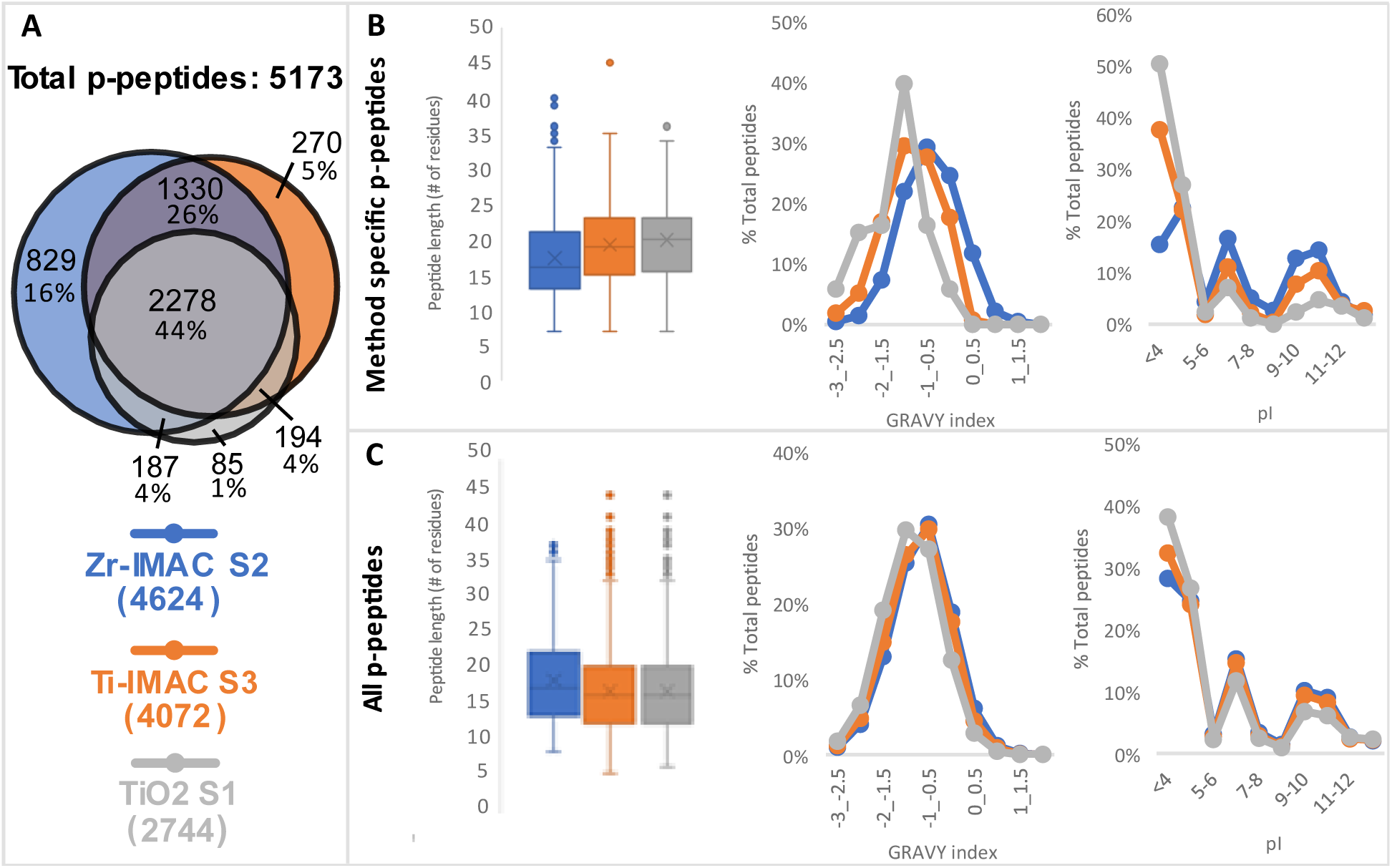
Feature analysis of phosphopeptides (p-pep) identified in the optimized methods. A) At the top, Venn diagram of the p-pep identified by each method (the number of p-pep and percentage relative to total). Combination of p-pep identified (5173). At the bottom, number of identified p-pep in each method: 4624 (S2/Zr-IMAC, blue), 4072 (S3/Ti-IMAC, orange) and 2744 (S1/ TiO_2_, grey). B) Feature analysis of enriched p-pep found only in one of the methods: S2/Zr-IMAC (829 p-pep, blue), S3/Ti-IMAC (270 p-pep, orange) and S1/ TiO_2_ (85 p-pep, grey). From left to right, boxplot of peptide length distribution with mean marked as an X, GRAVY indexes distribution and pI values distribution. Lower values of GRAVY index represent less hydrophobicity. C) Feature analysis of all enriched p-pep from S2/Zr-IMAC (4624 p-pep, blue), S3/Ti-IMAC (4072 p-pep, orange) and S1/ TiO_2_ (2744 p-pep, grey). From left to right, boxplot of peptide length distribution with mean marked as an X, GRAVY indexes distribution and pI values distribution.

To understand the binding properties of each bead type and their potential complementarity, the number of method specific p-pep identified (S2/Zr-IMAC 829, S3/Ti-IMAC 270, S1/ TiO_2_ 85, Figure 4A) were characterized according to their hydrophobicity, pI value and peptide length (Figure 4B). As a control, the same characteristics were analysed for all identified p-pep in each method (S2/Zr-IMAC 4624, S3/Ti-IMAC 4072, S1/ TiO_2_ 2744, figure 4C).

In figure 4B, the unique phosphopeptidesbound by Ti-IMAC and TiO_2_ were longer than those recovered by Zr-IMAC (median of 19, 20 and 16 amino acids (a.a.) respectively. Considering all phosphopeptides (Figure 4C), the length distribution of Ti-IMAC and TiO_2_ shifted to shorter sequences while Zr-IMAC maintained a similar profile as for peptides (medians of 17, 17 and 16 a.a. respectively).

The hydrophobicity curve of S2/Zr-IMAC was notably skewed to the right in Figure 4B and more subtly in Figure 4C, showing a higher affinity of Zr-IMAC for hydrophobic sequences compared to the other methods, whereas S1/ TiO_2_ was skewed more to the left, i.e. a more hydrophilic preference.

The pI values of each condition highlighted a greater proportion of acidic sequences in TiO_2_ (Figure 4B and 4C, right) as a result of the less acidic environment. On the other hand, Zr-IMAC and Ti-IMAC had more similar profiles, though Zr-IMAC showed a more distributed proportion of its unique p-pep.

The physical features of the uniquely found phosphopeptides from the standard and optimized solvents for each bead chemistry were also scrutinized (Supplementary Figure 4). More acidic and hydrophilic sequences bound to TiO_2_ when using the higher pH solvent S1 (0.1% Acetic acid) since the selectivity decreased and non-protonated carboxylated groups could interact with the metal oxide. This is also reflected by the decreased number of multiply phosphorylated peptides (Figure 3B) as a result of a more competitive environment. Changing 1M of Glycolic acid (Std) to either 0.1M of GA (S2) or 0.1M LA (S3) increased binding of more hydrophobic and basic peptides. This change in hydrophobicity could be affected by the number of phosphate groups in the peptides, as Std/Zr-IMAC unique peptides were mostly multi-phosphorylated while S2/Zr-IMAC had mostly mono-phosphorylated peptides (Supplementary Figure 5). Furthermore, the average peptide length was lower in the optimized conditions compared to the standard for all chemistries.

**Figure 5.**
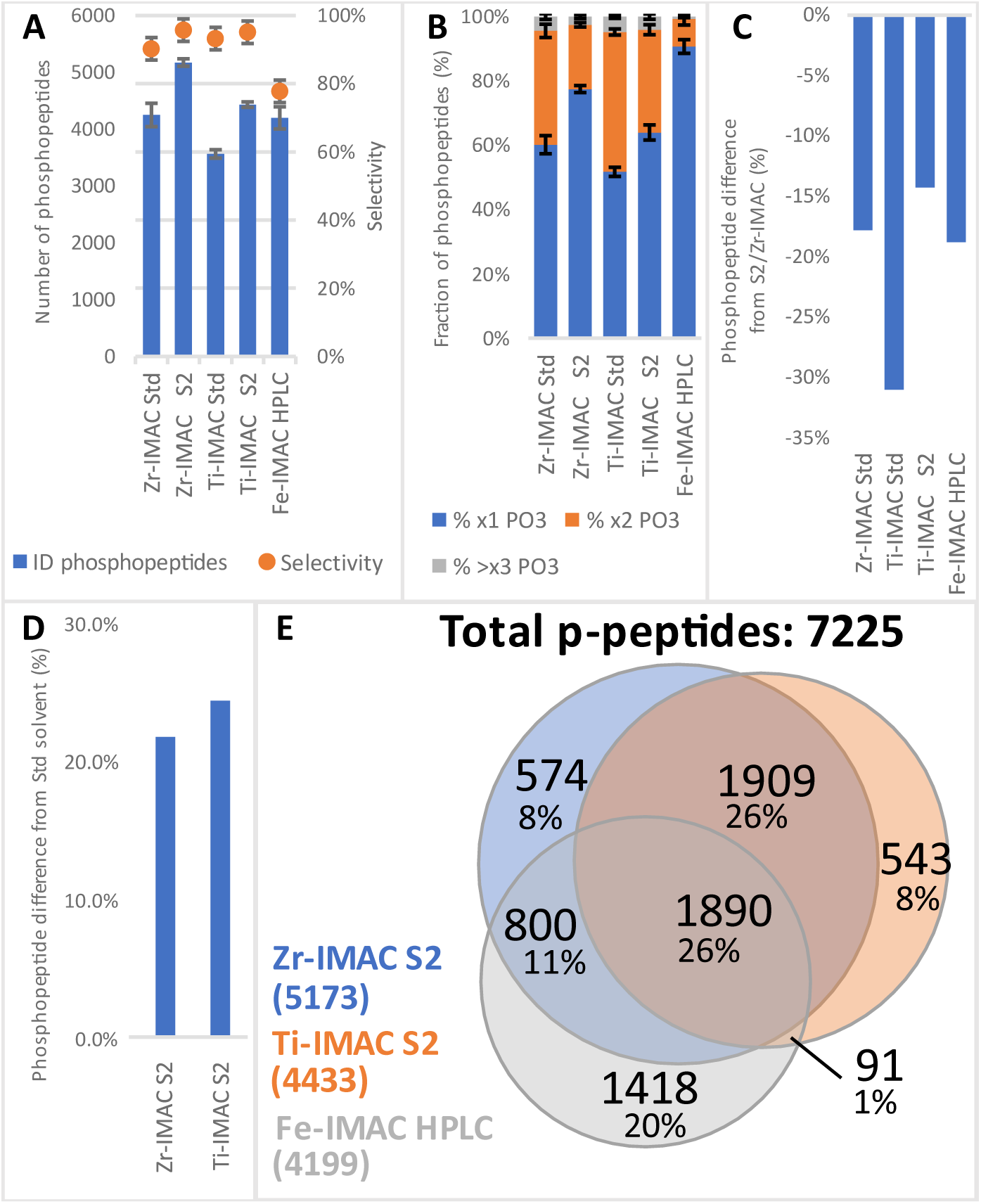
Phosphopeptide (p-pep) profiling of MagReSyn® microparticles by LC-MS/MS on Q Exactive HF. (A) Number of p-pep identified for each condition (blue bars) and the selectivity of each enrichment (orange dots). (B) Mono (blue), di (orange) and multiphosphorylated (grey) peptide distribution in each condition. (C) Differences in percentage of identified p-pep in each condition compared to the optimized S2/Zr-IMAC enrichment method. (D) Differences in percentage of identified p-pep between the optimized solvent (S2) and the standard (Std). (E) Venn diagram of identified p-pep from S2/Zr-IMAC (blue), S2/Ti-IMAC (orange) and Fe-IMAC (grey).

### Comparison of optimized microparticles and solvents with Fe-IMAC HPLC

Our previous results showed that Zr-IMAC recovered the highest number of phosphopeptides (4624) with the optimized solvent S2 (80% MeCN + 5% TFA + 0.1M GA). We hypothesised that S2 could improve Ti-IMAC microparticles performance on complex digests just as it did with Zr-IMAC. For this purpose, Zr-IMAC and Ti-IMAC magnetic microbeads were tested manually with both the standard (Std) and optimized (S2) conditions and compared with a well-known method for p-pep enrichment: Fe-IMAC (LC-format). Since Fe (III) ions have different chemical and coordination properties, like lower charge state and binding to just one supporting ligand (NTA)[51], we hypothesized that both magnetic materials would be more selective and capture higher numbers of p-pep than Fe-IMAC. The phosphopeptide enriched fractions were analysed by LC-MS/MS on a Q-Exactive HF mass spectrometer and the raw files were processed for peptide identification using Proteome Discoverer with Mascot as the search engine (Figure 1B, right branch).

The optimized enrichment with Zr-IMAC microparticles identified the highest number of phosphopeptides (5173), followed by S2/Ti-IMAC, Std/Zr-IMAC, Fe-IMAC HPLC and Std/Ti-IMAC (4433, 4249, 4199 and 3564, respectively, Figure 5A). All the enrichments performed with magnetic bead microparticles were highly selective towards p-pep with more than 90% of the total peptides assigned as phosphopeptides, while the selectivity of Fe-IMAC HPLC was significantly lower (78%) in our hands.

As expected, changing the binding solvent to S2 increased the number of single phosphorylated peptides in Zr-IMAC (78% x1 PO3), affecting Ti-IMAC as well (64% x1 PO3) to a higher extent than when using S3 (see Figure 3B and 5B). Moreover, Fe-IMAC mostly captured mono-phosphorylated peptides. Only 19% of the phosphopeptides carried more than one phosphate-group per peptide (Figure 5B), whereas Std Ti-IMAC and Std Zr-IMAC had a more even distribution (52% and 60% x1 PO3 respectively). Even though the distributions differ, the numbers of identified di-and multi-phosphorylated peptides was similar in Std/Zr-IMAC, Std/Ti-IMAC and S2/Ti-IMAC. They identified 1691, 1720 and 1597 multi-phosphorylated peptides respectively (Supplementary Figure 5).

Figure 5C shows the differences in bound phosphopeptides of all the conditions relative to S2/Zr-IMAC. Optimized Zr-IMAC was the best performing method in terms of total identifications (hence the negative values) binding 14% and 31% more p-pep than Ti-IMAC in S2 and Std solvents respectively, and 19% more than Fe-IMAC. Additionally, S2 improved the performance of Ti-IMAC with 24.4% more p-pep identified compared to Std/Ti-IMAC (Figure 5D), a greater difference than when solvent S3 was used (Figure 3C).

Collectively, all three methods contributed to the identification of a total of 7225 phosphopeptides (Figure 5E). Each method provided results and information that complemented the other methods. S2/Zr-IMAC and S2/Ti-IMAC identified altogether 5807 p-pep (80.4% of the total), complementing Zr-IMAC alone by 11% and Ti-IMAC alone by 24% (Figure 5E). In combination with Fe-IMAC, S2/Zr-IMAC identified 92.5% (6682) of the total number of identified p-pep, while Fe-IMAC alone covered 58.1% albeit with many method-unique phosphopeptides (1418). This opens the possibility for complementary enrichment, e.g. utilising Fe-IMAC for monophosphorylated peptide enrichment and Zr-IMAC for multiple phosphorylated peptides.

Examining the uniquely found phosphopeptides for each method (S2/Zr-IMAC: 574, S2/Ti-IMAC: 543, Fe-IMAC: 1418, Figure 5E), Fe-IMAC HPLC showed a preference towards more basic sequences than the other two methods, as it recovered peptides with higher pI values (Supplementary Figure 8). Zr- and Ti-IMAC showed similar profiles of acidity and basicity, though Ti-IMAC bound more acidic phosphopeptides than Zr-IMAC. Peptide hydrophobicity was higher in Zr-IMAC than Ti- and Fe-IMAC, and Fe-IMAC identified shorter peptides than Zr-IMAC and Ti-IMAC (Supplementary Figure 8).

Method complementarity was also observed at the individual phosphoprotein level. For example, phosphorylation sites in a single protein were localized by either S2/Zr-IMAC, S2/Ti-IMAC or Fe-IMAC. As an illustration, human protein scribble homolog (UniProt ac. No. Q14160), a scaffold protein found in the plasma membrane of cells involved in morphogenesis regulation. A total of 12 high confidence phosphorylation sites (phosphoRS > 95%) were localized in the protein. Four sites were found solely by S2/Zr-IMAC (S37, S853, S1348, S1508), one by S2/Ti-IMAC (S1306) and one by Fe-IMAC (S1448) (Supplementary Figure 7).

We investigated the differences in the bound phosphopeptides between binding solvents Std and S2 in Ti-IMAC and Zr-IMAC (Supplementary Figure 9), which presented very similar profiles to those of the previous experiments (Supplementary Figure 4). In this case, the optimized conditions recovered longer peptide sequences than the control, with Zr-IMAC having similar lengths and properties in both Std and S2. Ti-IMAC, on the other hand, showed more distinct differences, binding more acidic peptides with solvent S2 than with Std.

Finally, we looked at the number of identified phosphorylated proteins between the different methods and conditions (Supplementary Figure 10). Fe-IMAC HPLC identified the highest number of proteins compared to Zr- and Ti-IMAC in Std solvent (1891, 1682 and 1456 identifications, respectively). However, when using solvent S2 both Zr-IMAC and Ti-IMAC identified 17% more phosphoproteins each, with Zr-IMAC surpassing Fe-IMAC HPLC at 1965 identifications, again revealing the complementarity of these methods.

## DISCUSSION

Mass spectrometry-based phosphoproteomics is an important bioanalytical tool for investigating dynamic cell signalling processes and discover potential biomarkers for diseases, such as cancer and diabetes [14, 17, 23, 65]. Therefore, it is imperative to develop efficient, sensitive and automated methods for enrichment of phosphopeptides in fast and reproducible analytical and computational workflows. Towards this goal we optimized Zr-IMAC in a magnetic bead format and demonstrated that Zr-IMAC is a reliable tool for large-scale phosphoproteome analysis.

We first investigated six solvent compositions for three types of magnetic microparticles (Zr-IMAC, Ti-IMAC and TiO_2_), and we found ideal conditions for phosphopeptide enrichment for each one. Next, we compared each material under standard and optimized conditions using 200ug of HepG2/C3A cell protein digest. The improved Zr-IMAC enrichment captured the highest number of p-pep with ~97% selectivity, identifying 19% more p-pep than in standard conditions (Figures 3A, 3B and 3C). We then proceeded to test Zr-IMAC and Ti-IMAC magnetic microparticles against Fe-IMAC HPLC based p-pep enrichment [38, 66], using both control and the optimized solvents. Compared to Fe-IMAC, our results revealed high selectivity of all magnetic microbeads based methods, greater coverage of multiphosphorylated peptides and, in the case of optimized Zr-IMAC, higher numbers of p-pep and phosphorylated proteins (Figures 5A, 5B, Supplementary Figure 10). In short, Zr-IMAC demonstrated to be on par with more popular enrichment methods and outperforming them with some optimization of enrichment conditions. Using magnetic microparticles allowed for faster and parallel workflows for processing samples. We could perform 16 magnetic particle-based enrichment experiments in parallel in one hour using manual sample processing, and with application of a magnetic handling station up to 96 samples can be processed in parallel in less than one hour[31, 67, 68].

Reducing the concentration of hydroxy acid in the binding solvent increased the number of identified phosphopeptides for all tested affinity materials (Figure 3A and 3C). Since pH and ionic strength of the mobile phase affect peptide retention [69], a possible explanation could be that the high concentration of glycolic acid (1M) in the solvent increased competition with the phosphate groups of the peptides, while optimized solvents all had lower concentrations of acid (0.1M), possibly reducing the effect of competition. This improvement in p-pep recovery also affected phosphoprotein identification, where once again S2/Zr-IMAC outperformed the other condition (Supplementary Figure 2) without any significant differences in terms of protein localization in the cell (Supplementary Figure 3).

We observed a moderate increase in hydrophobic sequences when using the optimized solvents in Zr- and Ti-IMAC, as well as a large increase in acidic peptides for TiO_2_ (Figures 4B and 4C, Supplementary Figure 4). This change in peptide properties could be related to the decrease of proton and hydroxy acid concentrations in solution. Weaker ionic strength and less competition for chelation could improve the diffusion of highly hydrophobic sequences to the aqueous phase where the electrophile metals are located [34]. Consequently, in Std solvent (5% TFA, 1M GA) these peptides would not bind efficiently to the metal cations and would be lost during the washing steps, explaining also why we could see lower numbers of monophosphorylated peptides under these conditions, being less hydrophilic than multiphosphorylated peptides.

As solvent S2 proved to enhance Zr-IMAC enrichment, we also evaluated its effect on Ti-IMAC. We further compared these magnetic materials with Fe-IMAC HPLC, a well-established method for phosphopeptide enrichment [38, 70, 71]. Even though we identified more phosphorylated proteins in Fe-IMAC than in Std/Zr-IMAC and Std/Ti-IMAC, optimized Zr-IMAC enrichment outperformed both Fe-IMAC and Ti-IMAC, identifying more phosphoproteins and p-pep with an excellent selectivity of over 90% (Figure 5A, Supplemental figure 10). Moreover, the Fe-IMAC HPLC column revealed a bias towards monophosphorylated peptides and lower selectivity than the magnetic microparticles (Figures 5A and 5B), which, in contrast, they showed a more distributed binding of mono- and multi-phosphorylated peptides. Nevertheless, both Ti- and Zr-IMAC had a much higher preference to acidic sequences than Fe-IMAC. This behaviour could be explained by the strong interactivity of Ti4+ and Zr4+ with oxygen groups [52–54, 72], whereas the weaker Lewis acid Fe3+ could interact with other side-chain groups.

Combining results obtained by using Zr-IMAC magnetic microparticles with Fe-IMAC HPLC, we were able to boost our p-pep numbers by 21% and identify 15% more phosphoproteins (Figure 5E, Supplementary Figure 10). We also observed complementarity of these enrichment methods in mapping multiple phosphorylation sites in individual proteins (Supplementary Figure 7). Our study, like others, highlights the importance of applying several enrichment strategies for comprehensive phosphoprotein analysis, since potentially valuable information can be missed when using only one method.

In conclusion, we demonstrated that Zr-IMAC magnetic microparticles are generally applicable to phosphopeptide enrichment in phosphoproteomics workflows and it performs on par, if not better, than currently used IMAC and MOAC techniques. We captured high numbers of p-pep with optimized sample loading solvent for Zr-IMAC while maintaining excellent selectivity. The Zr-IMAC magnetic microbead format complements currently used phosphopeptide enrichment methods and is easily automated using robotic sample handling. We conclude that Zr-IMAC is widely applicable in functional phosphoproteomics studies and cell signalling research in biology and biomedicine.

## ABBREVIATIONS

AcA: Acetic acid
MeCN: Acetonitrile
ABC: Ammonium bicarbonate
DDA: Data-dependent adquisition
FA: Formic acid
GA: Glycolic acid
HCD: Higher-energy collisional dissociation
IMAC: Immobilized Metal ion Affinity Chromatography
IAA: Iodoacetamide
mgf: Mascot Generic File
MOAC: Metal Oxide Affinity Chromatography
p-pep: Phosphopeptides
SDC: Sodium deoxycholate
S1/S2/S3/S4/S5: Solvent 1/2/3/4/5
TA: Tartaric acid
Ti: Titanium
TiO_2_: Titanium oxide
TCEP: Tris(2-carboxyethyl)phosphine hydrochloride
Zr: Zirconium
CAA: 2-Chloroacetamide

## Acknowledgements

This study was part of the activities of the EU Horizon 2020 Innovative Training Network (ITN) *Biocapture* (Grant agreement 722171). We thank Adelina Rogowska-Wrzesinska and Helle Frandsen for providing hepatocyte cells. Proteomics and mass spectrometry research at SDU is supported by generous grants to the VILLUM Center for Bioanalytical Sciences (VILLUM Foundation grant no. 7292, to O.N.J.) and PRO-MS: Danish National Mass Spectrometry Platform for Functional Proteomics (grant no. 5072-00007B, to O.N.J.). The proteomics and mass spectrometry research at CSIR is supported by generous grants from the Council of Scientific and Industrial Research (2017/2018 Thematic Programme grant no. TP-04-2018).

## Disclaimer/Disclosure

Stoychev S. and Jordaan J. are employees of ReSyn Biosciences and have financial interests in the company. Govender I., and Naicker P., work on collaborative projects across CSIR and ReSyn Biosciences.

