## Supplemental figures for "Zirconium(IV)-IMAC for phosphopeptide enrichment in phosphoproteomics"

SUPPLEMENTARY FIGURE 1

SUPPLEMENTARY FIGURE 2

SUPPLEMENTARY FIGURE 3

SUPPLEMENTARY FIGURE 4

SUPPLEMENTARY FIGURE 5

SUPPLEMENTARY FIGURE 6

SUPPLEMENTARY FIGURE 7

SUPPLEMENTARY FIGURE 8

SUPPLEMENTARY FIGURE 9

SUPPLEMENTARY FIGURE 10

SUPPLEMENTARY FIGURE 1. List of isotopically labelled peptides and identified phosphopeptides from alpha- and beta-Casein.

VPQ[L_C13N15]EI[V_C13N15]PN[pSer]AEER. Ratio: 1. Mass: 1672 Da, [M+H]+: m/z 1673

YKVPQLE[I_C13N15]VPN[pSer]AEER. Ratio: 3. Mass: 1657 Da, [M+H]+: m/z 1658

FQ[pSer]EEQQQTEDE[L_C13N15]QDK. Ratio: 3. Mass: 2067 Da, [M+H]+: m/z 2068

| α-casein S1 (α-S1) and S2 (α-S2), and β-casein (β-C) | | | |
|---|---|---|---|
| **Sequence** | **Protein** | **# PO3** | **(M+H)+** |
| EQLSTSEENSK | (α-S2-(141–151)) | 2 | 1411.50 |
| EQLSTSEENSKK | (α-S2-(141–152)) | 2 | 1539.60 |
| TVDMESTEVFTK | (α-S2-(153–164)) | 1 | 1466.61 |
| TVDMESTEVFTKK | (α-S2-(153–165)) | 1 | 1594.70 |
| VPQLEIVPNSAEER | (α-S1-(121–134)) | 1 | 1660.79 |
| YLGYLEIVPNSAEER | (α-S1)b | 1 | 1832.83 |
| DIGSESTEDQAMEDIK | (α-S1-(58–73)) | 2 | 1927.69 |
| YKVPQLEIVPNSAEER | (α-S1-(119–134)) | 1 | 1951.95 |
| FQSEEQQQTEDELQDK | (β-C-(33–48)) | 1 | 2061.83 |
| NMAINPSKENLCSTFCK | (α-S2-(39-55)) | 1 | 2093.80 |

SUPPLEMENTARY FIGURE 2. Venn diagrams of identified phosphorylated proteins in Zr-IMAC, Ti-IMAC and TiO2 magnetic microspheres under two different binding solvents: control (Std) and optimized (S1, S2, S3).

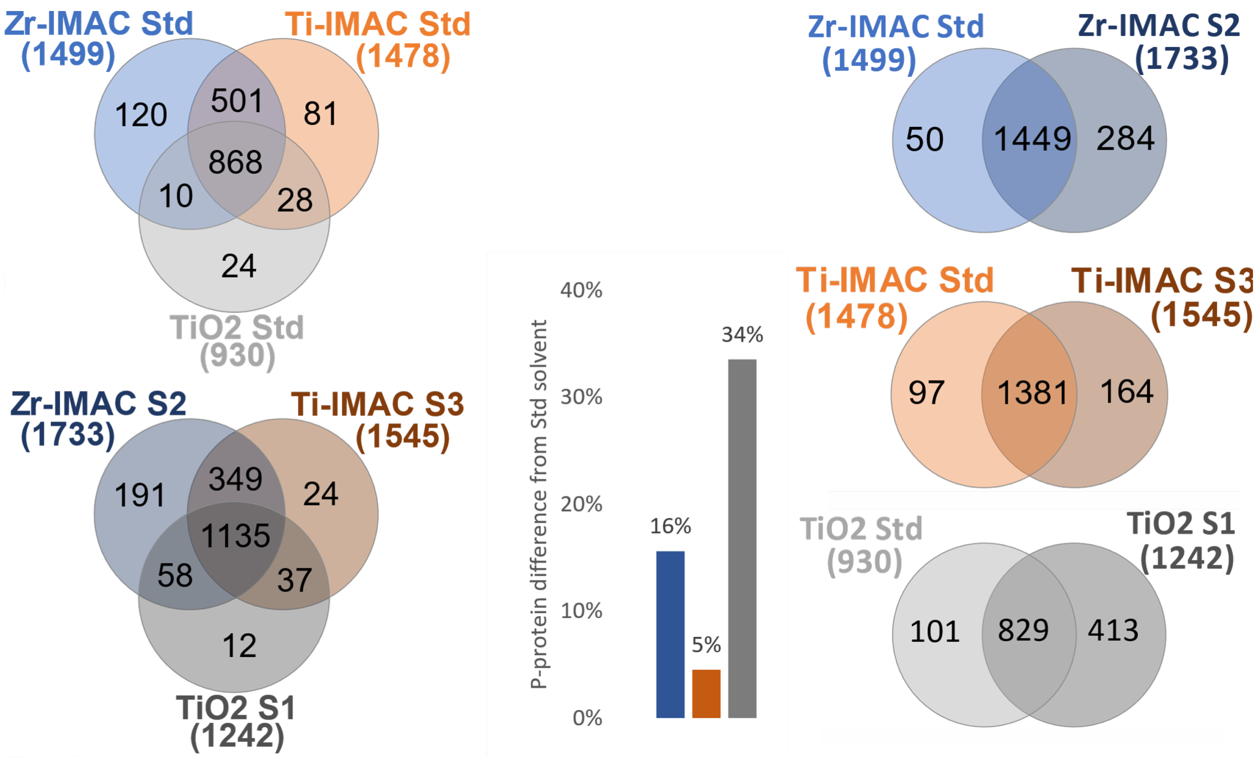

SUPPLEMENTARY FIGURE 3. Cellular component GO analysis of identified phosphorylated proteins in optimized methods S2/Zr-IMAC, S3/Ti-IMAC and S1/TiO2.

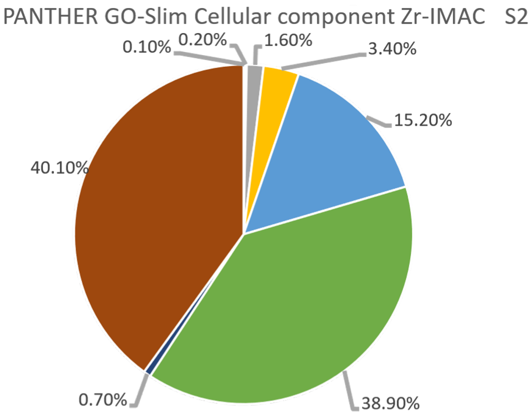


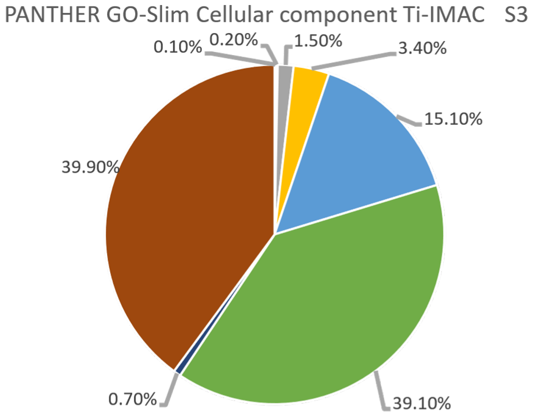


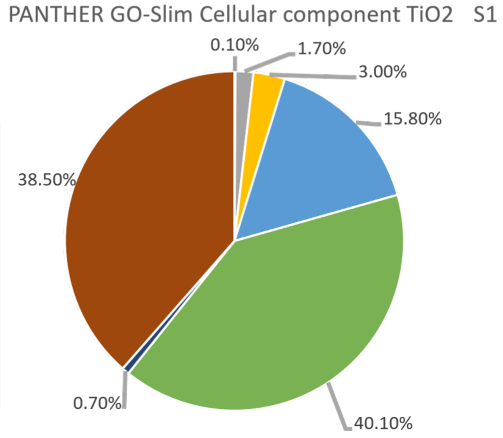


| Cellular component | Zr-IMAC S2 | Ti-IMAC S3 | TiO2 S1 |
|---|---|---|---|
| Synapse (GO:0045202) | 0.10% | 0.10% | 0% |
| Supramolecular complex (GO:0099080) | 0.20% | 0.20% | 0.10% |
| Cell junction (GO:0030054) | 1.60% | 1.50% | 1.70% |
| Membrane (GO:0016020) | 3.40% | 3.40% | 3.00% |
| Protein-containing complex (GO:0032991) | 15.20% | 15.10% | 15.80% |
| Organelle (GO:0043226) | 38.90% | 39.10% | 40.10% |
| Extracellular region (GO:0005576) | 0.70% | 0.70% | 0.70% |
| Cell (GO:0005623) | 40.10% | 39.90% | 38.50% |

SUPPLEMENTARY FIGURE 4. Feature analysis of uniquely enriched phosphopeptides from binding solvent Std (darker colour) and optimized (lighter colour) in three bead chemistries: Zr-IMAC (top, blue), Ti-IMAC (middle, orange) and TiO2 (bottom, grey). Plots represent the distribution of the total peptides in that condition for, from left to right, GRAVY Index, pI values and peptide length. Lower values of GRAVY index represent less hydrophobicity. Venn diagram (right) shows the peptides identified between the solvent conditions in each chemistry.

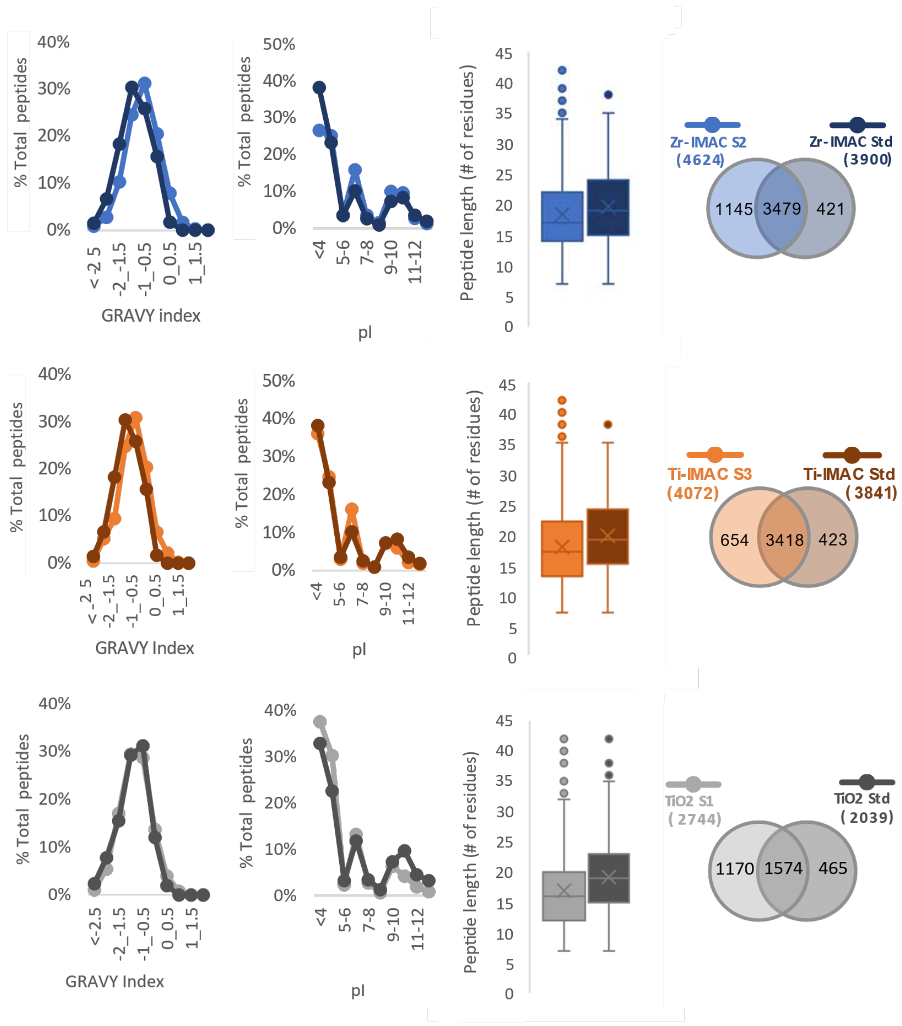

SUPPLEMENTARY FIGURE 5. GRAVY index profiles of unique mono- and multi-phosphorylated peptides of Zr-IMAC in solvent Std and S2.

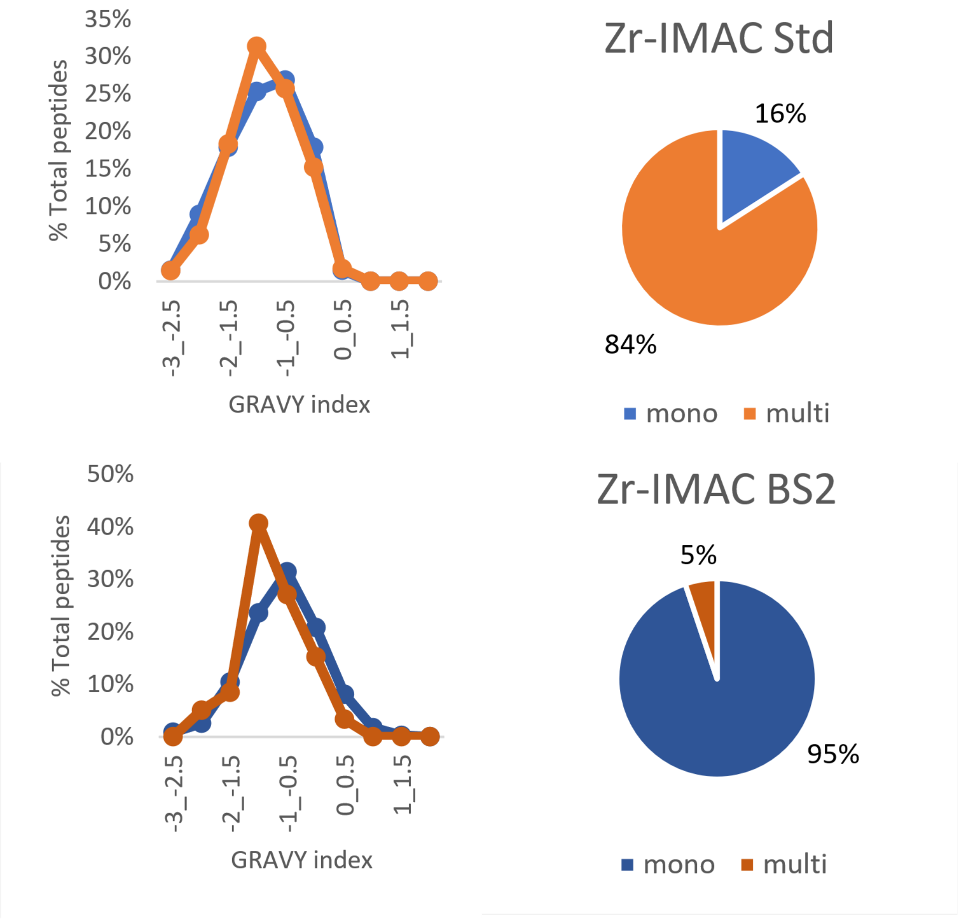

SUPPLEMENTARY FIGURE 6. Number of each phosphopetide types identified in all experiments.

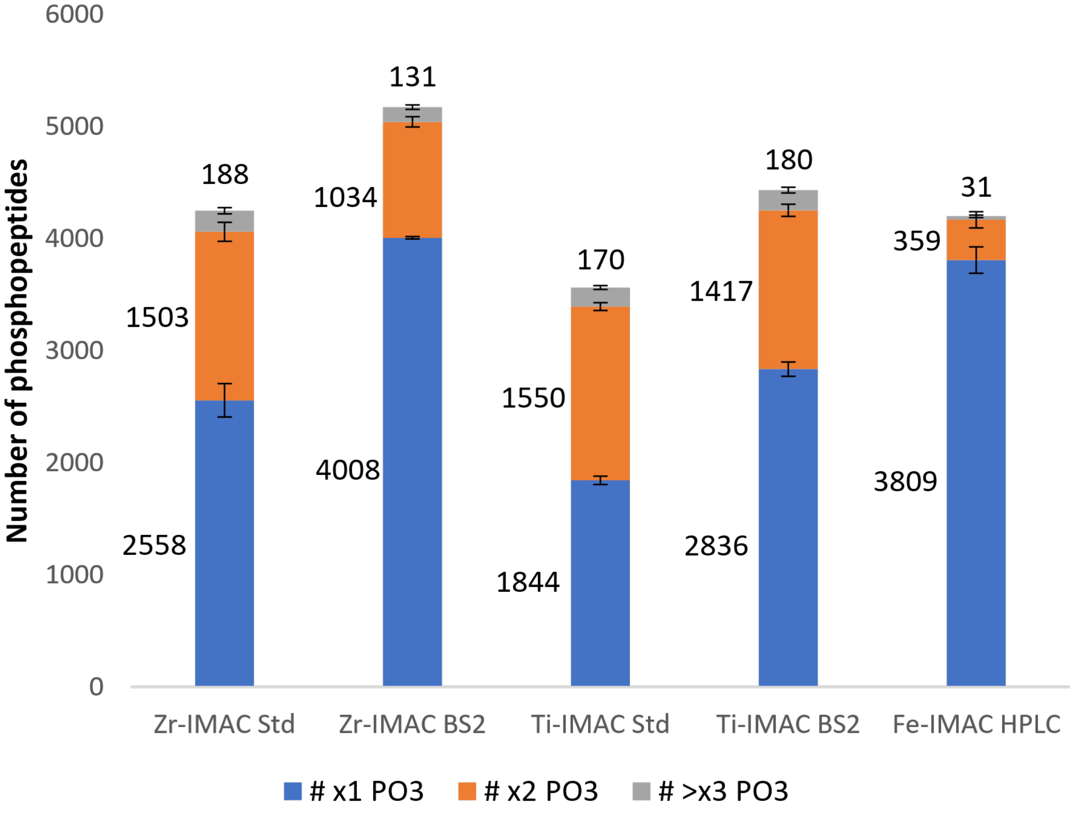

SUPPLEMENTARY FIGURE 7. Amino acid sequence of Protein scribble homolog (UniProt ac. No. Q14160). The method specific phosphorylation sites are colour coded: S2/Zr-IMAC (green), S2/Ti-IMAC (yellow), Fe-IMAC HPLC (blue). Identified in two methods: Zr-IMAC and Ti-IMAC (purple), Zr-IMAC and Fe-IMAC HPLC (orange). Identifed in all methods (red). Underneath, table of identified phosphopeptides from Q14160 in either S2/Zr-IMAC, S2/Ti-IMAC or Fe-IMAC.

Sequence: [Q14160] Protein scribble homolog

```
MLKCIPLWRCNRHVESVDKRHCSLQAVPEEIYRYSRSLEELLLDANQLRELPKPFFRLLNLRKLGLSDNEIQRLPPEVAN   80
FMQLVELDVSRNDIPEIPESIKFCKALEIADFSGNPLSRLPDGFTQLRSLAHLALNDVSLQALPGDVGNLANLVTLELRE  160
NLLKSLPASLSFLVKLEQLDLGGNDLEVLPDTLGALPNLRELWLDRNQLSALPPELGNLRRLVCLDVSENRLEELPAELG  240
GLVLLTDLLLSQNLLRRLPDGIGQLKQLSILKVDQNRLCEVTEAIGDCENLSELILTENLLMALPRSLGKLTKLTNLNVD  320
RNHLEALPPEIGGCVALSVLSLRDNRLAVLPPELAHTTELHVLDVAGNRLQSLPFALTHLNLKALWLAENQAQPMLRFQT  400
EDDARTGEKVLTCYLLPQQPPPSLEDAGQQGSLSETWSDAPPSRVSVIQFLEAPIGDEDAEEAAAEKRGLQRRATPHPSE  480
LKVMKRSIEGRRSEACPCQPDSGSPLPAEEEKRLSAESGLSEDSRPSASTVSEAEPEGPSAEAQGGSQQEATTAGGEEDA  560
EEDYQEPTVHFAEDALLPGDDREIEEGQPEAPWTLPGGRQRLIRKDTPHYKKHFKISKLPQPEAVVALLQGMQPDGEGPV  640
APGGWHNGPHAPWAPRAQKEEEEEEEGSPQEEEVEEEEENRAEEEEASTEEEDKEGAVVSAPSVKGVSFDQANNLLIEPA  720
RIEEEELTLTILRQTGGLGISIAGGKGSTPYKGDDEGIFISRVSEEGPAARAGVRVGDKLLEVNGVALQGAEHHEAVEAL  800
RGAGTAVQMRVWRERMVEPENAVTITPLRPEDDYSPRERRGGGLRLPLLPPESPGPLRQRHVACLARSERGLGFSIAGGK  880
GSTPYRAGDAGIFVSRIAEGGAAHRAGTLQVGDRVLSINGVDVTEARHDHAVSLLTAASPTIALLLEREAGGPLPPSPLP  960
HSSPPTAAVATTSITTATPGVPGLPSLAPSLLAAALEGPYPVEEIRLPRAGGPLGLSIVGGSDHSSHPFGVQEPGVFISK1040
VLPRGLAARSGLRVGDRILAVNGQDVRDATHQEAVSALLRPCLELSLLVRRDPAPPGLRELCIQKAPGERLGISIRGGAR1120
GHAGNPRDPTDEGIFISKVSPTGAAGRDGRLRVGLRLLEVNQQSLLGLTHGEAVQLLRSVGDTLTVLVCDGFEASTDAAL1200
EVSPGVIANPFAAGIGHRNSLESISSIDRELSPEGPGKEKELPGQTLHWGPEATEAAGRGLQPLKLDYRALAAVPSAGSV1280
QRVPSGAAGGKMAESPCSPSGQQPPSPPSPDELPANVKQAYRAFAAVPTSHPPEDAPAQPPTPGPAASPEQLSFRERQKY1360
FELEVRVPQAEGPPKRVSLVGADDLRKMQEEEARKLQQKRAQMLREAAEAGAEARLALDGETLGEEEQEDEQPPWASPSP1440
TSRQSPASPPPLGGGAPVRTAKAERRHQERLRVQSPEPPAPERALSPAELRALEAEKRALWRAARMKSLEQDALRAQMVL1520
SRSQEGRGTRGPLERLAEAPSPAPTPSPTPVEDLGPQTSTSPGRLSPDFAEELRSLEPSPSPGPQEEDGEVALVLLGRPS1600
PGAVGPEDVALCSSRRPVRPGRRGLGPVPS
```

**Zr-IMAC S2** **Ti-IMAC S2** **Fe-IMAC HPLC**

**Zr-/Ti-/Fe- IMAC** **Zr-/Ti- IMAC** **Zr-/Fe- IMAC**

★ **Not found in UniProt**

| Sequence | Master Protein Accessions | Modifications in Proteins | Confidence Zr-IMAC S2 | Confidence Ti-IMAC S2 | Confidence Fe-IMAC |
|---|---|---|---|---|---|
| RVSLVGADDLRK | Q14160 | 1xPhospho [S1378(100)] | High | Not Found | High |
| LPLLPPESPGPLR | Q14160 | 1xPhospho [S853(100)] | High | Peak Found | Not Found |
| NSLESISSIDR | Q14160 | 1xPhospho [S1220(100)] | High | High | High |
| MAESPCSPSGQQPPSPPSPDELPANVK | Q14160 | 2xPhospho [S1306(100); S1309(100)] | Peak Found | High | Not Found |
| MAESPCSPSGQQPPSPPSPDELPANVK | Q14160 | 1xPhospho [S1309(100)] | High | High | Not Found |
| MVEPENAVTITPLRPEDDYSPR | Q14160 | 1xPhospho [S835(100)] | High | High | Not Found |
| MKSLEQDALR | Q14160 | 1xPhospho [S1508(100)] | High | Not Found | Peak Found |
| SLEELLLDANQLR * | Q14160 | 1xPhospho [S37(100)] | High | Peak Found | Not Found |
| AFAAVPTSHPPEDAPAQPPTPGPAASPEQLSFR | Q14160 | 1xPhospho [S1348(100)] | High | Not Found | Not Found |
| LAEAPSPAPTPSPTPVEDLGPQTSTSPGRLSPDFAEELR | Q14160 | 2xPhospho [S1547(97); S1566(100)] | High | High | Not Found |
| NSLESISSIDRELSPEGPGK | Q14160 | 2xPhospho [S1220(100); S] | Not Found | High | Not Found |
| RVSLVGADDLR | Q14160 | 1xPhospho [S1378(100)] | High | Not Found | High |
| LALDGETLGEEEQEDEQPPWASPSPTSR | Q14160 | 1xPhospho [S1439(99.1)] | High | High | Not Found |
| QSPASPPPLGGGAPVR | Q14160 | 1xPhospho [S1448(100)] | Not Found | Not Found | High |

* Found in Q14160 and Q9BTT6

SUPPLEMENTARY FIGURE 8. Feature analysis of uniquely identified phosphopeptides in Zr-IMAC, Ti-IMAC and Fe-IMAC, based on peptide length, hydrophobicity and pI value. A venn diagram shows the nuber of shared and unique phosphopeptides between the methods.

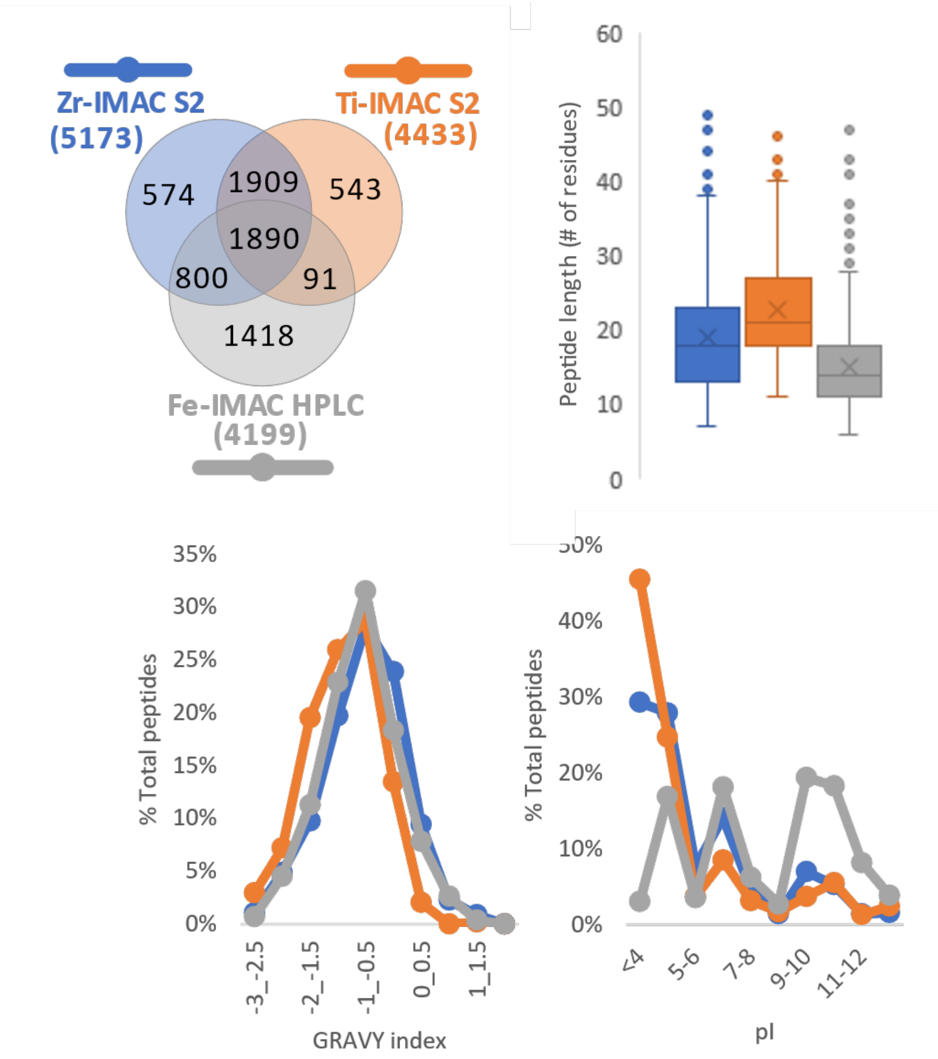

SUPPLEMENTARY FIGURE 9. Peptide feature analysis of Zr-IMAC and Ti-IMAC in Std and S2 solvents.

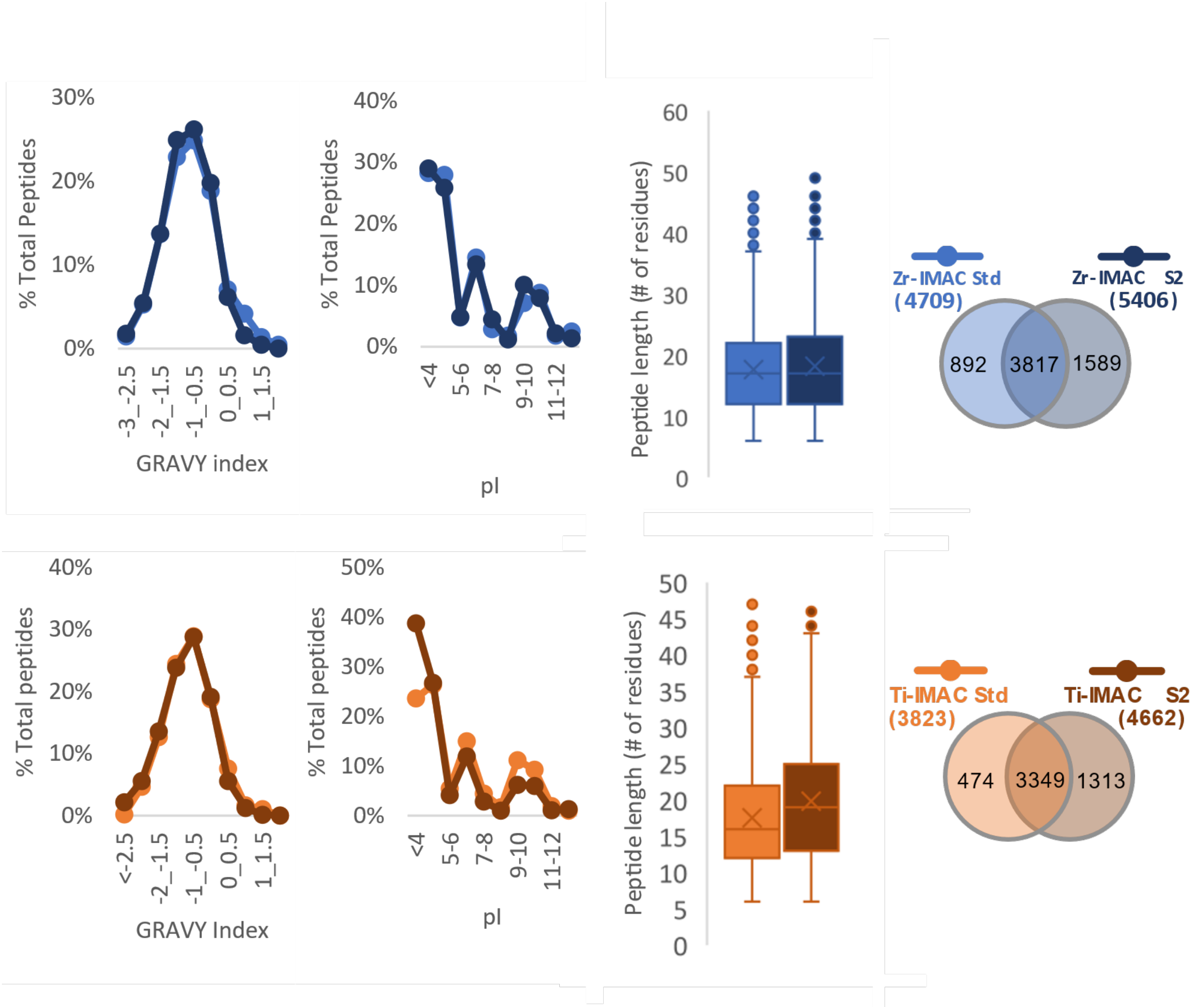

SUPPLEMENTARY FIGURE 10. Venn diagrams of identified phosphorylated proteins in Zr-IMAC and Ti-IMAC magnetic microspheres and Fe-IMAC HPLC under two different binding solvents: Std and S2.

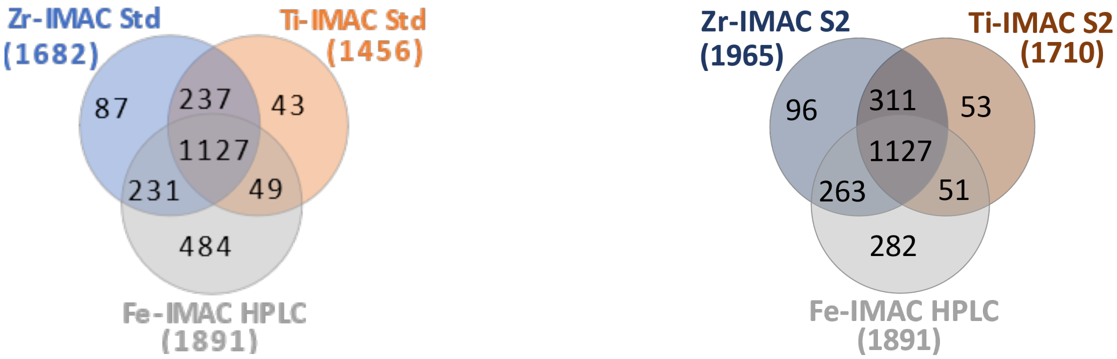
